# Structure of the extracellular region of the bacterial type VIIb secretion system subunit EsaA

**DOI:** 10.1101/2020.08.17.254201

**Authors:** Timothy A. Klein, Dirk W. Grebenc, Shil Y. Gandhi, Vraj S. Shah, Youngchang Kim, John C. Whitney

**Affiliations:** Michael DeGroote Institute for Infectious Disease Research, McMaster University, Hamilton, ON, L8S 4K1, Canada; Department of Biochemistry and Biomedical Sciences, McMaster University, Hamilton, ON, L8S 4K1, Canada; Structural Biology Center, X-ray Science, Argonne National Laboratory, Argonne, Illinois, USA; David Braley Centre for Antibiotic Discovery, McMaster University, Hamilton, ON, L8S 4K1, Canada

## Abstract

Gram-positive bacteria use type VII secretion systems (T7SSs) to export effector proteins that manipulate the physiology of nearby prokaryotic and eukaryotic cells. Several mycobacterial T7SSs have established roles in virulence. By contrast, recent work has demonstrated that the genetically distinct T7SSb pathway found in Firmicutes bacteria more often functions to mediate interbacterial competition. A lack of structural information on the T7SSb has limited the understanding of effector export by this protein secretion apparatus. In this work, we present the 2.4Å crystal structure of the extracellular region of the elusive T7SSb subunit EsaA from *Streptococcus gallolyticus*. Our structure reveals that homodimeric EsaA is an elongated, arrow-shaped protein with a surface-accessible ‘tip’, which serves as a receptor for lytic bacteriophages in some species of bacteria. Because it is the only T7SSb subunit large enough to traverse the thick peptidoglycan layer of Firmicutes bacteria, we propose that EsaA plays a critical role in transporting effectors across the entirety of the Gram-positive cell envelope.

## Introduction

Protein secretion is a critical aspect of bacterial physiology and requires the use of membrane-embedded secretion apparatuses. In addition to the general secretory pathway and the twin-arginine translocase, many species of Gram-positive bacteria use type VII secretion systems (T7SSs) for protein export (Abdallah et al., 2007). T7SSs are used by bacteria belonging to the phyla Actinobacteria and Firmicutes and are divided into T7SSa and T7SSb. This distinction reflects differences in T7SS subunit composition between these two distantly related groups of Gram-positive bacteria (Klein et al., 2020). The T7SSa was originally discovered in *Mycobacterium tuberculosis* where it acts as a virulence factor that facilitates immune evasion and phagosomal escape during infection, whereas the T7SSb was initially characterized in *Staphylococcus aureus* and has been shown to play a dual role in pathogenesis and interbacterial competition (Cao et al., 2016; Gao et al., 2004; Ohr et al., 2017; Ulhuq et al., 2020). The interkingdom-targeting capability of the T7SSb has also been demonstrated in the opportunistic pathogen *Streptococcus intermedius* with the antibacterial activity being attributed to the NAD^+^ hydrolase effector TelB and the cell wall precursor degrading effector TelC (Hasegawa et al., 2017; Klein et al., 2018; Whitney et al., 2017). The T7SSb pathways of *Bacillus subtilis* and *Enterococcus faecalis* were also recently shown to antagonize competitor bacteria (Tassinari et al., 2020; Chatterjee et al., 2020).

Much of our current understanding of the T7SS has resulted from studies on effector function, which can often explain the phenotypes associated with a given T7SS pathway. Less well understood is the mechanism of T7SS effector export across the cell envelope. Recent structural analyses have begun to elucidate the ultrastructure of T7SS apparatuses and provide clues as to how this secretion apparatus facilitates protein export (Famelis et al., 2019; Poweleit et al., 2019; Rosenberg et al., 2015). However, these studies have largely focused on T7SSa apparatuses. Of the four major structural proteins that make up the T7SSa, only the EccC/EssC/YukB ATPase is conserved in T7SSb systems. The other three T7SSa subunits, EccB, EccD, and EccE, possess no sequence homology to the EssA, EssB, and EsaA components of the T7SSb and consequently, the two systems likely form distinct structures that may not share a common mechanism for protein export.

EsaA is perhaps the least understood of the T7SSb structural components. Transposon mutagenesis in *S. aureus* initially suggested that EsaA was dispensable for effector secretion but subsequent characterization showed that this subunit is likely essential for T7SSb-dependent protein export (Burts et al., 2005; Kneuper et al., 2014). No structural data exists for EsaA, but analysis of its membrane topology suggests it consists of a large soluble region flanked by N- and C-terminal transmembrane domains (TMDs) (Ahmed et al., 2018; Mietrach et al., 2019). Proteomic analyses of intact *S. aureus* cells has shown that EsaA is surface exposed and that its soluble domain may extend into the extracellular milieu (Dreisbach et al., 2010). Furthermore, studies in *B. subtilis* have shown that the EsaA homologue YueB is the cell surface receptor for the SPP1 bacteriophage (Sao-Jose et al., 2004; Sao-Jose et al., 2006). Similarly, many strains of *E. faecalis* possess the EsaA paralogue <u>P</u>hage Infection <u>P</u>rotein (PIP), which serves as a receptor for Enterococcal phage (Duerkop et al., 2016). The prediction that EsaA extends from the plasma membrane to the cell surface make it unique among the T7SSb subunits because the other structural proteins have either extracellular domains that are too small to span the estimated 30-50 nm thick peptidoglycan layer of Firmicutes bacteria or are entirely intracellular (Tassinari et al., 2020; Vollmer et al., 2008).

In this study, we present the first crystal structure of the extracellular domain of EsaA, revealing a novel protein fold characterized by a highly elongated, arrow-shaped homodimer comprised of three distinct domains. Using cysteine cross-linking, we show that EsaA dimers occur *in vivo* and propose that upon multimerization with the other subunits of the T7SSb, form a conduit that facilitates effector export across the cell envelope of Gram-positive bacteria.

## Results

### EsaA is required for the secretion of EsxA and Tel effector proteins from *S. intermedius*

Given the conflicting reports on the essentiality of EsaA for T7S, we first examined the consequences of inactivating *esaA* on effector export using the model T7SSb bacterium *S. intermedius*. Characterized T7SSb systems export two major families of effectors: small, alpha-helical WXG100 proteins whose precise function is unknown; and large, multi-domain LXG proteins that possess C-terminal toxin domains. *S. intermedius* strain B196 exports a single WXG100 effector, EsxA, and the three LXG effectors TelA, TelB and TelC (Whitney et al., 2017). Consistent with functioning as a core structural subunit of the T7SSb apparatus, we found that replacement of the *esaA* gene with a kanamycin resistance cassette yielded a *S. intermedius* strain that was no longer able to export detectable levels of EsxA and TelC into culture supernatants (Figure 1A). Similarly, supernatant NADase activity, which is indicative of TelB secretion, was reduced to levels comparable to that of a T7SSb-inactivated strain, Δ*essC*. (Figure 1B). Importantly, we found that export of EsxA and TelC, as well as TelB-dependent NADase activity could be restored by plasmid-based expression of EsaA indicating that our allelic replacement approach did not affect the expression of genes encoding other structural subunits of the T7SSb, which are part of a five-gene cluster that also contains *esaA*. Together, these data indicate that *esaA* is required for WXG100 and LXG effector export in *S. intermedius*.

**Figure 1.**
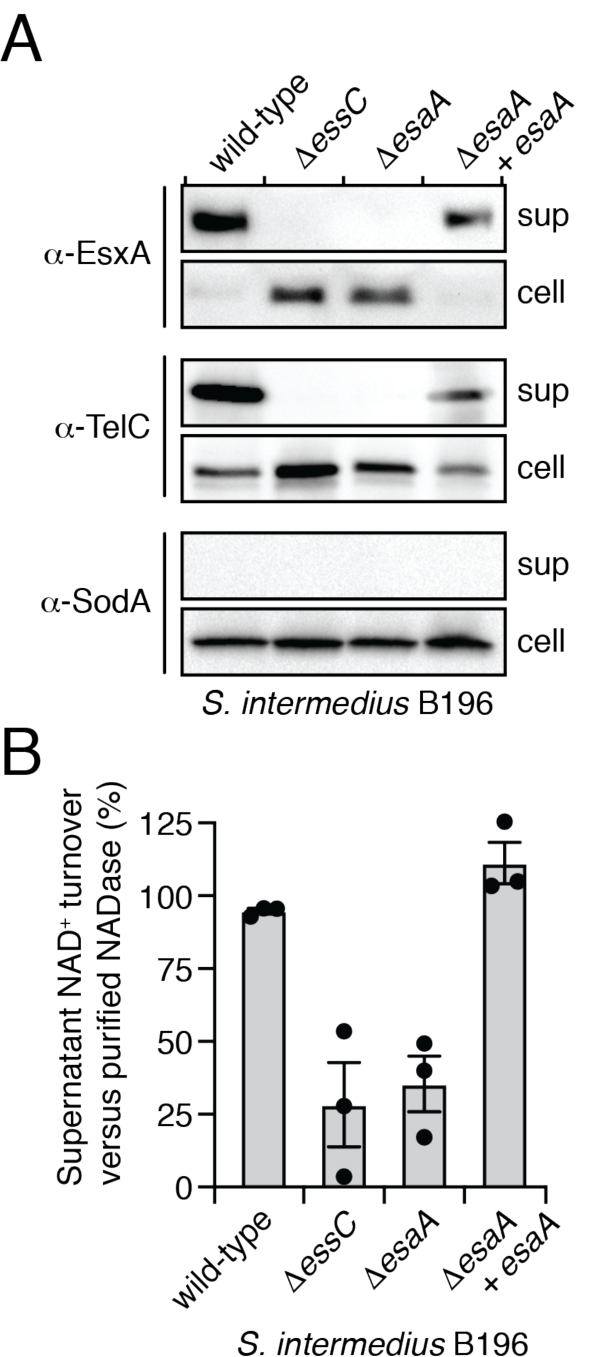
EsaA is required for WXG-100 and LXG effector export by *S. intermedius*. (A) Western blot analysis of the cell and supernatant fractions of the indicated *S. intermedius* B196 strains. EsxA and TelC belong to the WXG-100 and LXG families of T7SSb effectors, respectively. The Δ*essC* strain is used as a secretion deficient control. Superoxide dismutase A (SodA) is used as a cell lysis control. (B) Supernatant NADase activity, indicative of T7SSb-dependent TelB secretion, in cultures of the indicated *S. intermedius* B196 strains. This assay was done in triplicate and all values were calculated as a fraction of NAD^+^ turnover compared to purified NADase Tse6 (Whitney et al., 2015). The data displayed represent three independent replicates. Error bars reflect standard error of the mean (SEM).

### Topology mapping of EsaA reveals a large extracellular domain

We next sought to examine the membrane topology of *S. intermedius* EsaA (*Si*EsaA). Though cell surface proteomics conducted on *S. aureus* suggest that the soluble region of EsaA exists extracellularly, this assertion has not been tested directly for any T7SSb-containing bacterium. Furthermore, the number of putative TMDs differs among EsaA homologues with *Si*EsaA having a single predicted TMD on either side of its soluble region whereas EsaA proteins from *S. aureus, E. faecalis, B. subtilis, Bacillus cereus* and *Listeria monocytogenes* possess five TMDs at their C-terminus (Figure S1).

After confirming that *Si*EsaA localizes to the membrane fraction of lysed *S. intermedius* cells (Figure 2A), we introduced a series of cysteine point mutations spaced approximately 150 amino acids apart within *Si*EsaA to map its membrane topology using a cysteine-reactive maleimide-conjugated fluorophore (Figure 2B). Plasmid-borne expression of each EsaA cysteine mutant in our *esaA* deletion strain restored T7SSb-dependent export of TelC, demonstrating that these mutations do not significantly affect EsaA function (Figure 2C). *Si*EsaA contains a single native cysteine residue predicted to reside in its N-terminal TMD, and we found that with intact cells this residue was inaccessible to the cysteine-reactive dye when analyzed by SDS-PAGE (Figure 2D). Similarly, *Si*EsaA variants harboring cysteine mutations near the N- (V8C) or C-terminus (F909C) of the protein did not react with the dye. By contrast, we found that cells expressing *Si*EsaA bearing V150C, F302C, S454C or S605C mutations, all of which reside within the predicted soluble region, yielded a prominent fluorescent band at the expected molecular weight of *Si*EsaA (Figure 2D). A fluorescent band absent in the wild-type control was also present in the V762C variant; however, this band migrates at a higher molecular weight than *Si*EsaA making it difficult to interpret. Collectively, our data indicate that *Si*EsaA is a membrane protein with a large extracellular domain and intracellular N- and C-termini.

**Figure 2.**
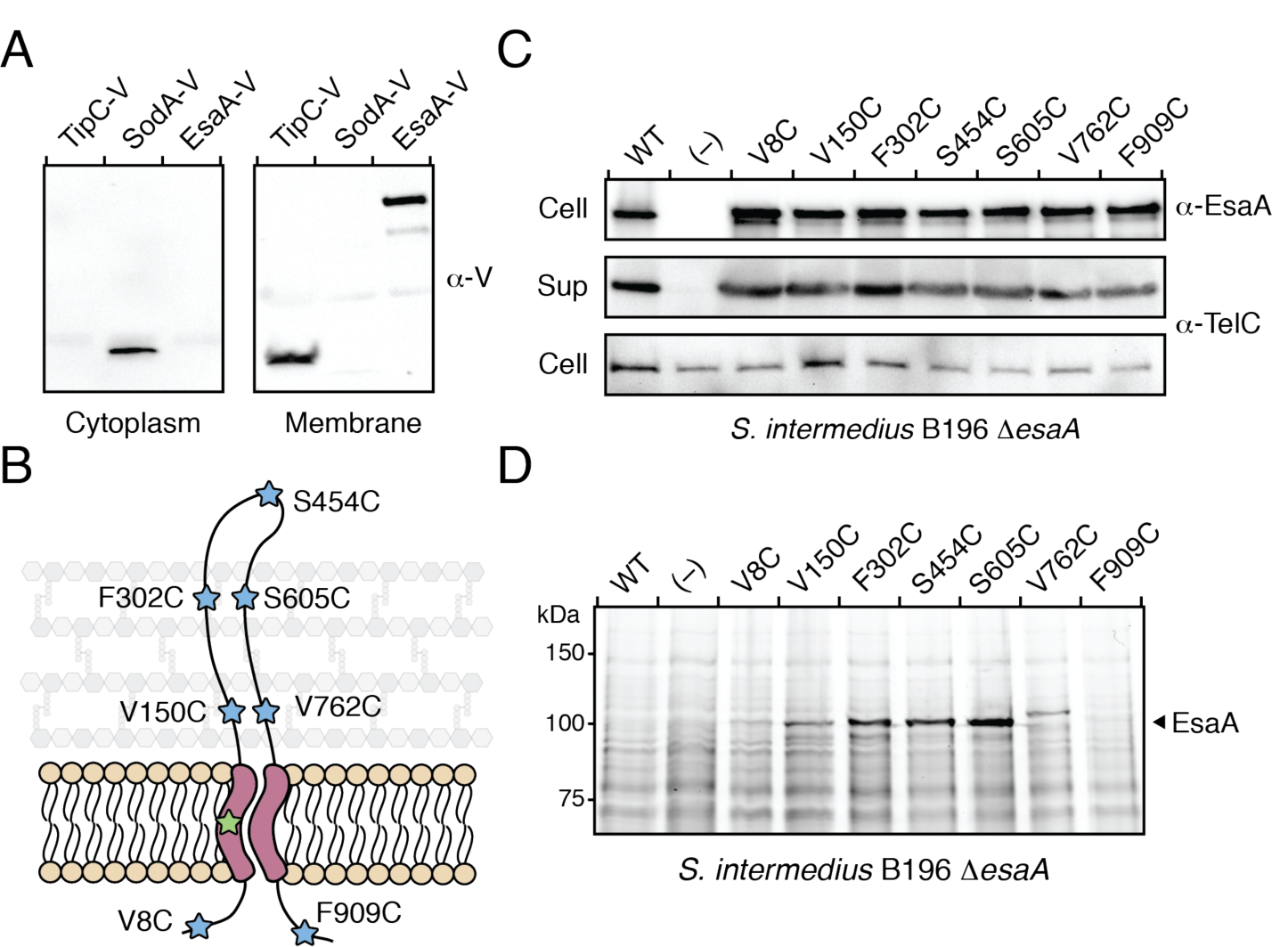
EsaA possesses a large extracellular domain. (A) EsaA fractionates with *S. intermedius* membranes. TipC and SodA are used as membrane and cytoplasmic controls, respectively. Proteins contain a C-terminal VSV-G tag and were detected by western blot using an α-VSV-G (α-V) primary antibody. (B) Predicted EsaA membrane topology depicting the location of each cysteine substitution site. Green star denotes the native cysteine residue present in EsaA whereas blue stars indicate cysteine mutations generated for topology mapping. (C) EsaA cysteine mutants are expressed and secrete TelC at levels similar to wild-type *S. intermedius*. (D) Cysteine mutations in the predicted extracellular domain of EsaA are accessible to a cysteine-reactive maleimide dye but those located near the N- and C-termini are not. EsaA migrates slightly above the 100kDa marker as indicated.

### Structure determination of an extracellular fragment of EsaA

Having mapped the membrane topology of *Si*EsaA, we next initiated structural studies on the large extracellular fragment of the protein to gain more insight into its function. Although we could readily express and purify a truncation of *Si*EsaA encompassing its entire extracellular region (residues 41-871), this protein fragment had a propensity to degrade. To identify a stable fragment of *Si*EsaA that would be more amenable to crystallization, we performed limited proteolysis with chymotrypsin and isolated a protease-resistant species spanning residues 234-790 (Figure S2). This fragment of *Si*EsaA crystallized readily but despite extensive optimization efforts, diffraction quality crystals could not be obtained. Using the boundary information obtained from our proteolysis experiments, we next tried a homologous EsaA fragment from *Streptococcus gallolyticus* ATCC 43143 (*Sg*EsaA_235-829_), which has 42.9% pairwise sequence identity to the equivalent region of *Si*EsaA (Figure 3A). Purified *Sg*EsaA_235-829_ formed diffraction quality crystals and the 2.4Å structure of SgEsaA_235-829_ was determined using selenium-incorporated protein and the single-wavelength anomalous dispersion technique (Table 1). Interestingly, the resulting electron density map only yielded interpretable density for a model encompassing residues 330-727 with an unmodeled gap from amino acids 514-554, suggesting that large portions of EsaA are disordered in the crystal lattice. The final model was refined to a *R*_work_/*R*_free_ of 0.21 and 0.26, respectively.

**Table 1:**
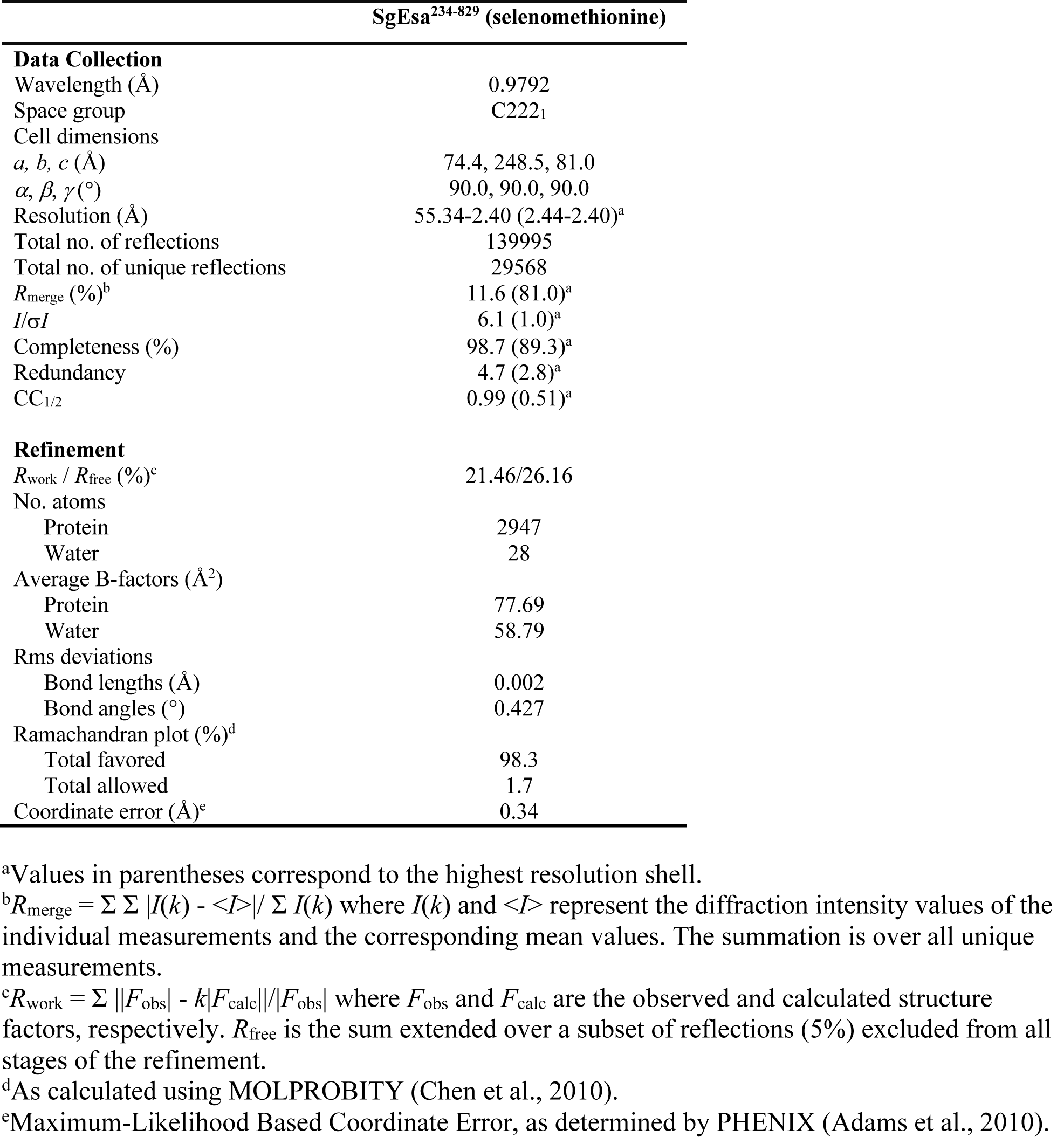
X-ray data collection and refinement statistics.

**Figure 3.**
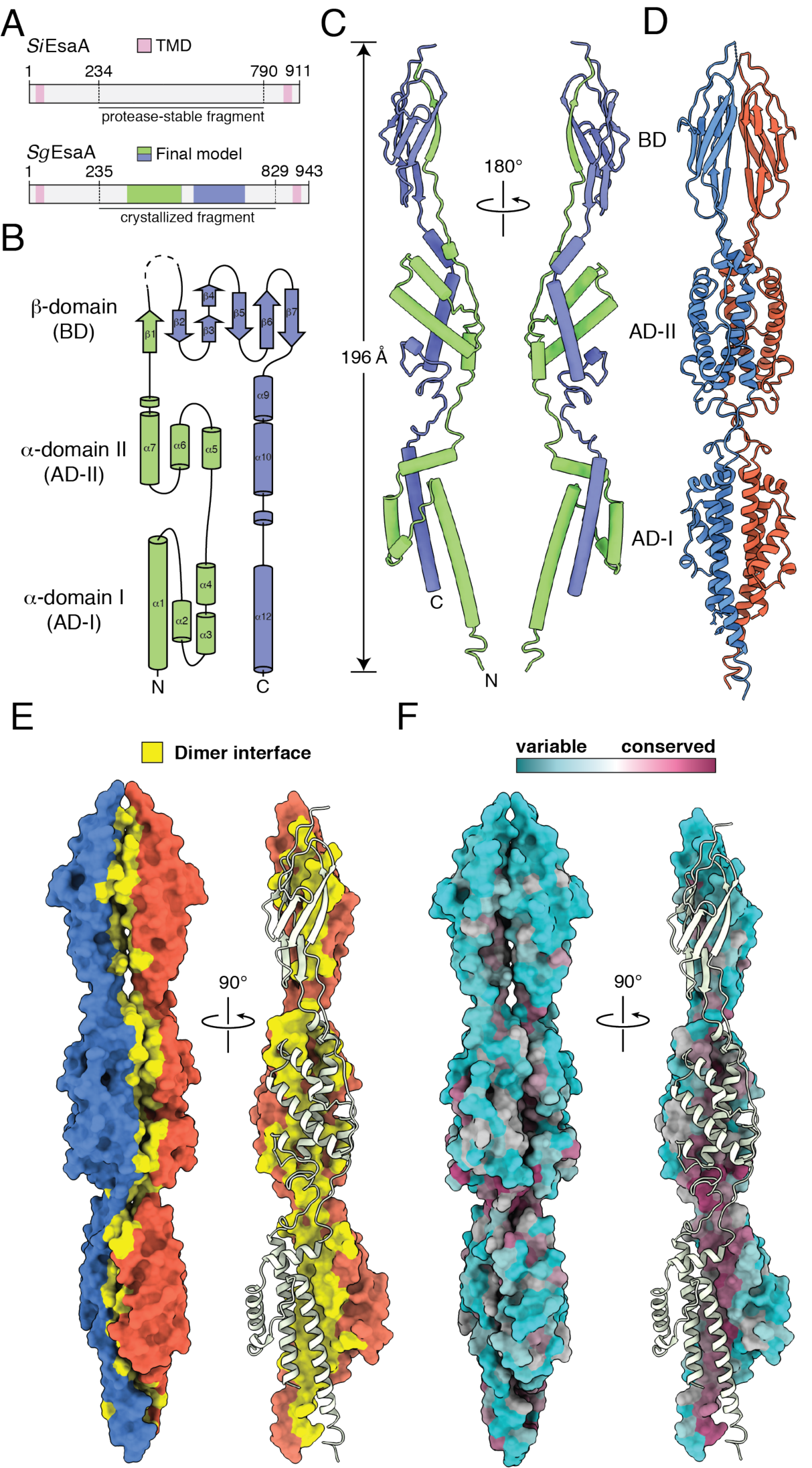
The extracellular domain of *Sg*EsaA adopts an arrow-shaped structure. (A) Domain architecture of *S. intermedius* B196 EsaA (*Si*EsaA) and *S. gallolyticus* ATCC 43143 EsaA (*Sg*EsaA) depicting the chymotrypsin-stable fragment of *Si*EsaA, the crystallized fragment of *Sg*EsaA and the regions of *Sg*EsaA for which interpretable electron density was observed in the crystal structure. (B) Topology diagram depicting the secondary structure elements comprising *Sg*EsaA_235-829_. Blue and green coloring is used to illustrate the ‘there and back again’ topology of the protein. (C) Pipes and planks model of *Sg*EsaA_235-829_ shown from two opposing views. Alpha helices and beta strands are denoted by tubes and arrows, respectively. The N- and C-termini are depicted on the left-hand model. (D) *Sg*EsaA_235-829_ dimers form an elongated structure. Red and blue ribbon coloring is used to differentiate each monomer within the dimer. (E) Surface representation of an *Sg*EsaA_235-829_ dimer shown from orthogonal viewpoints. Yellow coloring is used to highlight the buried surface area between *Sg*EsaA_235-829_ protomers. (F) Surface representation of an *Sg*EsaA_235-829_ dimer depicting residue-specific sequence conservation among EsaA homologous proteins. Details of the sequences used for conservation analysis can be found in Experimental Procedures. Model was generated using the ConSurf server (Ashkenazy et al., 2016).

### EsaA forms an elongated, arrow-shaped dimer

*Sg*EsaA_235-829_ forms a highly elongated structure comprised of two <u>a</u>lpha helical <u>d</u>omains (AD-I and AD-II) and a <u>b</u>eta-sheet <u>d</u>omain (BD) (Figures 3B and 3C). The modelled fragment adopts a ‘there and back again’ topology whereby the first half of *Sg*EsaA_235-829_ contributes secondary structure elements to each of the three domains over a linear distance of 196Å. Following a 180° turn that occurs within the unmodelled region between the β1 and β2 strands of the beta-sheet domain, the C-terminal half of the protein similarly contributes secondary structure to each domain with the C-terminus being located ∼20Å away from the N-terminus at the same pole (Figure 3B). In this arrangement, both the N- and C-terminal TMDs present in full-length EsaA would be connected to the alpha helical AD-I domain. Given the orientation of the termini, the directionality of the beta strands flanking the central unmodelled region, and the number of unmodelled amino acids in our structure, it is likely that the length of the entire extracellular region of EsaA is well in excess of the ∼200Å measured for our model. This finding provides a molecular explanation for how this protein is potentially able to traverse the approximately 30-50nm thick cell wall of Firmicutes bacteria (Vollmer et al., 2008).

Another striking feature of *Sg*EsaA_235-829_ is that it adopts a head-to-head, belly-to-belly homodimer that gives the protein its arrow-shaped appearance (Figure 3D). In this configuration, all three domains and the intervening connecting regions contribute to the dimerization interface (Figure 3E). Analysis of the dimer interface using the PDBePISA webserver indicates that dimer formation is highly favorable (Δ_i_G: -61.8kcal/mol) and generates 4436Å^2^ of buried surface area (Krissinel and Henrick, 2007). Mapping EsaA sequence conservation onto our structure reveals that the residues comprising the surface of EsaA are highly variable whereas the amino acids involved in homodimerization show a much higher level of conservation (Figure 3F). The amino acids lining the dimer interface are a mixture of hydrophobic, polar and acidic residues with tyrosine, leucine, threonine and glutamate being the most abundant. We speculate that the large surface area of the dimer interface combined with the abundance of hydrophobic residues participating in homodimerization indicates that EsaA likely exists as an obligate homodimer because solvent exposure of this surface in aqueous environments would bear a large entropic cost.

A comparison of *Sg*EsaA_235-829_ to previously determined structures in the Protein Data Bank using DALILITE revealed that the overall structure of *Sg*EsaA_235-829_ does not resemble proteins of known structure (Holm, 2020). The top hit from this search was the BID domain of the type IV secretion system (T4SS) effector protein Bep9 from *Bartonella clarridgeiae* (Z-score, 8.5; Cα root mean square deviation of 3.5Å over 100 aligned residues), which only shares structural similarity with AD-I of EsaA (Figure S3)(Stanger et al., 2017). BID domains comprise one part of a bipartite signal sequence found in some T4SS effectors and thus appear unrelated in terms of function. Based on these analyses, we conclude that EsaA adopts a novel protein fold.

### EsaA exists as dimer *in vitro* and *in vivo*

To test the biological significance of the EsaA homodimer observed in our crystal structure, we examined a truncation of *Sg*EsaA_235-829_ that more accurately reflects the modeled boundaries of our structure (*Sg*EsaA_332-725_) as well as the equivalent fragment of *Si*EsaA (*Si*EsaA_328-685_) by size exclusion chromatography coupled to multi-angle laser light scattering (SEC-MALS). SEC-MALS allows for the accurate determination of protein molecular mass in solution and therefore helps identify potentially artefactual oligomeric states induced by protein crystallization. For both proteins, the major peak yielded a molecular mass consistent with dimer formation and no evidence of EsaA monomers was observed in either case (Figure 4A and 4B). The SEC-MALS analysis of *Sg*EsaA_332-725_ also revealed the presence of high molecular weight aggregates but due to their heterogeneous nature and absence in the *Si*EsaA_328-685_ sample, we concluded that they likely do not represent biologically relevant assemblies of EsaA. In sum, the extracellular fragment of EsaA exists as a dimer in solution.

**Figure 4.**
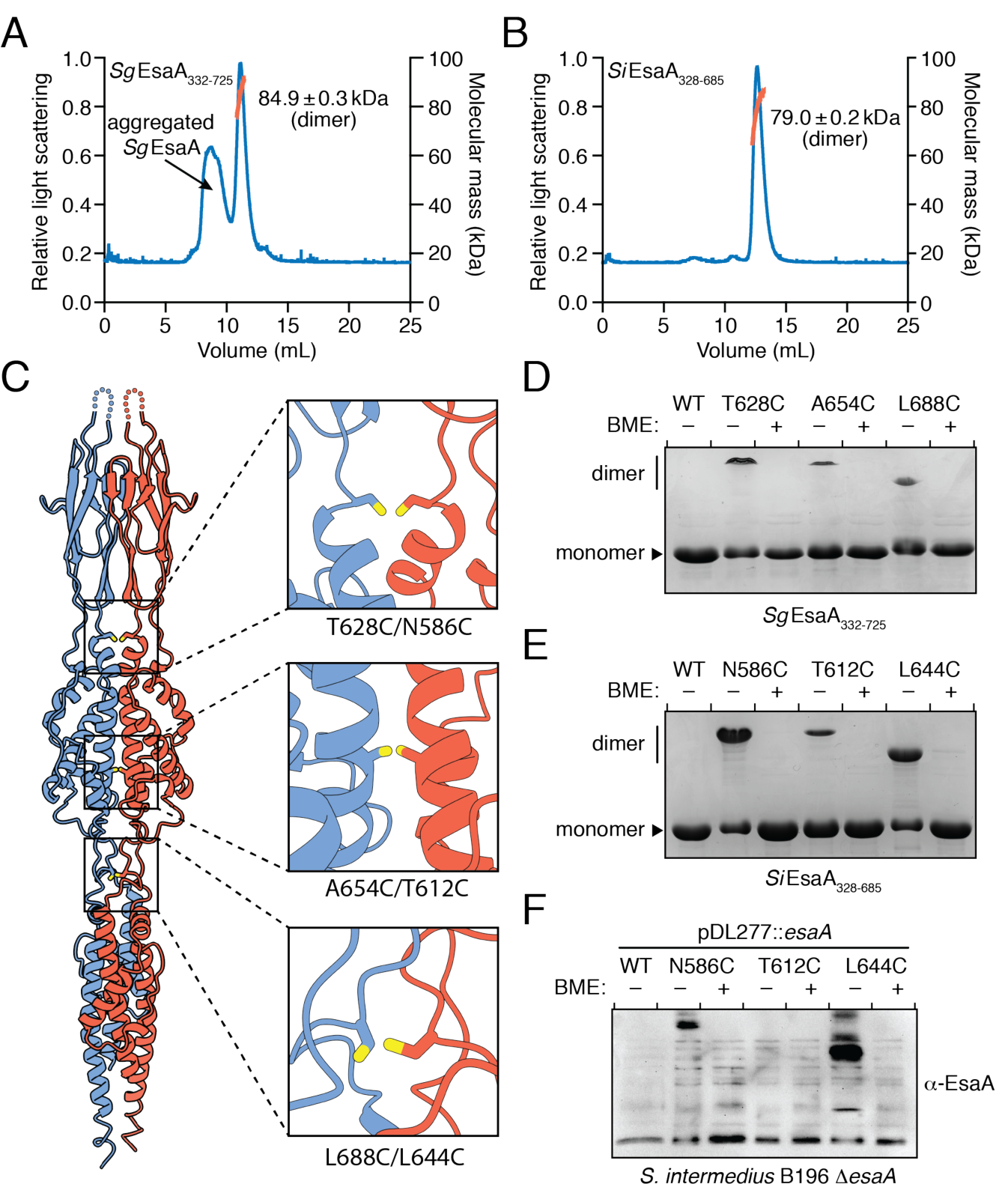
EsaA forms dimers *in vitro* and *in vivo*. (A-B) SEC-MALS analysis of *Sg*EsaA_332-725_ (A) and *Si*EsaA_328-685_ (B). Relative light scattering is plotted in blue and molecular weight is plotted in orange. The calculated molecular weights of the dimer peaks for both proteins are indicated. (C) Structure of *Sg*EsaA_332-725_ depicting the cysteine mutations chosen for cross-linking experiments. *Sg*EsaA_332-725_ protomers are depicted as blue and red ribbons with the hypothetical cysteine mutations shown as sticks. The identities of the residues normally found in these positions are indicated for both *Sg*EsaA (left) and *Si*EsaA (right). (D-E) Coomassie blue-stained gel demonstrating cysteine crosslinking for each of the purified *Sg*EsaA_332-725_ (D) and *Si*EsaA_328-685_ (E) cysteine variants. (F) Western blot analysis of *S. intermedius* B196 Δ*esaA* strains expressing wild-type EsaA or each of the indicated EsaA cysteine variants. BME, β-mercaptoethanol.

We next wanted to examine if EsaA dimerizes *in vivo* in a manner that is consistent with our crystal structure. To accomplish this, we inspected our *Sg*EsaA_235-829_ structure for amino acid residues within the dimer interface that would be expected to crosslink if mutated to cysteine. This analysis led to the identification of Thr628, found in in the linker region between the BD and AD-II, Ala654, located within the AD-II, and Leu688, which exists in the linker region between AD-I and AD-II (Figure 4C). We mutated each of these residues, along with the equivalent residues in *Si*EsaA (Asn586, Thr612 and Leu644), to cysteine and examined the ability of these variants to form covalent dimers. In support of the dimeric arrangement observed in our crystal structure, all six variants formed β-mercaptoethanol (BME)-sensitive crosslinks when the purified proteins were examined by SDS-PAGE (Figure 4D and 4E). Furthermore, when we introduced the *Si*EsaA cysteine variants into our *S. intermedius* B196 *esaA* deletion strain, BME-sensitive cysteine cross-links were observed in cells expressing either EsaA^N586C^ or EsaA^L644C^ (Figure 4F). Collectively, our cross-linking data suggest that the structure of *Si*EsaA is likely very similar to that of *Sg*EsaA in terms of overall fold and dimeric arrangement, and that dimeric EsaA represents a biologically relevant form of the protein.

### The structure of EsaA predicts the putative binding site for a bacteriophage receptor

EsaA homologous proteins are not only involved in type VII secretion but have also been shown to function as receptors for lytic bacteriophages (Sao-Jose et al., 2004). A recent analysis of Enterococcal phages identified a 160 amino acid hypervariable region within the EsaA homologous protein PIP (<u>P</u>hage <u>I</u>nfection <u>P</u>rotein) responsible for phage tropism among *E. faecalis* strains (Duerkop et al., 2016). The topology of EsaA combined with the domain organization revealed by our *Si*EsaA_328-685_ crystal structure suggest that the beta-sheet domain of this protein family is likely the surface exposed region, leading us to speculate that this region of the protein likely serves as the receptor for infecting phage. Indeed, mapping the hypervariable region of PIP proteins onto an EsaA-derived homology model of a representative PIP protein from *E. faecalis* V583 indicates that the phage tropism determining region identified by Duerkop et al. likely exists within the beta domain of EsaA homologous proteins (Figure S4).

## Discussion

Our structure of the extracellular region of EsaA has revealed the unique architecture of this enigmatic T7SSb subunit. EsaA adopts a novel protein fold that forms highly stable dimers *in vitro* and *in vivo*. The observation that T7SSb subunits form dimers is not without precedent as a recently determined crystal structure of full-length YukC (EssB) from *B. subtilis* found that this T7SSb subunit similarly homodimerizes (Tassinari et al., 2020). EssB/YukC also physically interacts with EsaA/YueB suggesting that these two proteins likely function together to facilitate protein secretion across the cell envelope (Ahmed et al., 2018).

Though T7SS structural components form dimers in crystals, current evidence indicates that the ultrastructure of an assembled T7SS apparatus involves hexamerization of the apparatus components. For example, the ESX-5 T7SSa from *Mycobacterium xenopi* exhibits six-fold symmetry and is proposed to contain 1:1:1:1 stoichiometry of the four T7SSa apparatus components EccB, EccC, EccD and EccE based on a 13Å negative stain electron microscopy (EM) map (Beckham et al., 2017). More recently, higher resolution cryo-EM structures of the ESX-3 T7SSa from *Mycobacterium smegmatis* have suggested a 1:1:2:1 protomer stoichiometry in which two EccD subunits interact with one subunit each of EccB, EccC, and EccE (Famelis et al., 2019; Poweleit et al., 2019). Though they are not homologues, EccB and EsaA are speculated to be functionally equivalent subunits between T7SSa and T7SSb pathways because they both possess large extracellular domains. However, our structure shows that both the overall structure and dimerization mode of EsaA is substantially different from that of EccB indicating that these T7SS subunits may have distinct functions. Furthermore, the extracellular region of EsaA is cell surface exposed whereas EccB predominantly exists in the mycobacterial periplasm. This observation suggests that additional factors may be involved in T7SSa-dependent effector export across the mycomembrane such as the EspB protein or members of the proline-glutamate and proline-proline-glutamate families of proteins (Solomonson et al., 2015; Wang et al., 2020). Ultimately, the structure of an intact T7SSb will be needed for an in-depth comparison between these intriguing protein export machines.

The *S. intermedius* T7SSb antibacterial effector TelC exerts toxicity in the inner wall zone (IWZ) by degrading the cell wall precursor lipid II present in the outer leaflet of the plasma membrane (Whitney et al., 2017). We previously used this unique site of action to provide evidence that the T7SSb exports effectors across the plasma membrane and the cell wall in a manner that bypasses the IWZ during transport (Klein et al., 2018). It is now apparent that EsaA, as the only T7SSb apparatus protein with an extended extracellular domain, may well form the conduit that allows for such transport. One of the defining characteristics of Gram-positive Firmicutes bacteria is the 30-50 nm thick peptidoglycan layer, which would likely prevent the diffusion of large LXG effectors from the IWZ to the milieu (Vollmer et al., 2008). Our structure of EsaA is 20 nm long and represents only a portion of the full-length protein. It is therefore within reason that EsaA extends across the entire cell wall to facilitate effector export from the cell. These observations, coupled with the abovementioned propensity for T7SS subunits to adopt six-point symmetry, lead to the tantalizing notion that EsaA dimers might trimerize to form a hexameric tube-shaped assembly. Such a structure would not only enable effector export from T7SSb-containing bacteria but may also facilitate the delivery of effectors into target cells.

## Experimental Procedures

### Bacterial strains, plasmids and growth conditions

All *S. intermedius* strains were generated from the *S. intermedius* B196 wild-type background. *E. coli* XL-1 Blue was used for plasmid maintenance. *E. coli* BL21 (DE3) CodonPlus and B834 (DE3) were used for the expression of methionine and selenomethionine containing proteins, respectively. Genomic DNA isolated from *S. intermedius* B196 and *S. gallolyticus* ATCC 43143 was used for cloning *Si*EsaA and *Sg*EsaA, respectively. A complete list of bacterial strains can be found in Table S1. pET29b and pDL277-derived plasmids were used for protein expression in *E. coli* and *S. intermedius*, respectively. pET29b-derived plasmids were generated by restriction enzyme-based cloning using the NdeI and XhoI restriction endonucleases and T4 DNA ligase. All constructs lacked their native stop codon resulting in the fusion of a vector encoded C-terminal his_6_-tag to facilitate protein purification after expression in *E. coli*. Cloning into pDL277 was performed similarly except with the BamHI and SalI restriction endonucleases. Additionally, the P96 promoter sequence from *Streptococcus pneumoniae* was fused upstream of all genes of interest using <u>s</u>plicing by <u>o</u>verlap <u>e</u>xtension (SOE) PCR to allow for gene expression in *S. intermedius* (Lo Sapio et al., 2012). All cysteine point mutations were generated by SOE PCR followed by restriction-enzyme based cloning into either pET29b or pDL277 with the abovementioned enzymes. A complete list of plasmids can be found in Table S2. All *E. coli* strains were grown overnight in lysogeny broth at 37°C at 225 rpm in a shaking incubator. Kanamycin (50 μg/mL) was added to the growth media for strains containing pET29b plasmids. All *S. intermedius* strains were grown in Todd Hewitt Broth supplemented with 0.5% yeast extract (THY) in a 37°C stationary 5% CO_2_ incubator. To ensure uniform growth rate, all *S. intermedius* strains were grown first on THY agar plates for 1-2 days prior to growth in THY broth. Strains harboring pDL277-derived plasmids were grown in media supplemented with spectinomycin (50μg/mL for *S. intermedius* or 100μg/mL for *E. coli*).

### DNA manipulation

*S. intermedius* and *S. gallolyticus* genomic DNA was prepared by resuspending cell pellets in InstaGene Matrix (Bio-Rad). Primers were synthesized by Integrated DNA Technology (IDT). Molecular cloning was performed using Q5 polymerase, restriction enzymes, and T4 DNA ligase from New England Biolabs (NEB). Sanger sequencing was performed by Genewiz Incorporated.

### Transformation of *S. intermedius*

*S. intermedius* transformation with either plasmid or linear DNA were performed as previously described (Tomoyasu et al., 2010). In short, overnight cultures were back diluted 1:10 into 2 ml THY broth supplemented with 3 μL of 10 mg/ml *S. intermedius* competence stimulating peptide (DSRIRMGFDFSKLFGK, synthesized by Genscript) and incubated at 37°C, 5% CO_2_ for 2 hours. Approximately 100-500ng of plasmid, or linear insert DNA was added and cultures were briefly vortexed before incubating for another 3 hours. 100 μl of culture was then plated on the appropriate selective media (either 50 μg/ml spectinomycin, 250 μg/ml kanamycin, or both).

### Gene deletion in *S. intermedius* by allelic replacement

SOE PCR was used generate a pETduet-1 plasmid containing the *kanR* cassette from the pBAV1k-e plasmid under the control of the spectinomycin promoter from pDL277. The spectinomycin promoter-kanamycin resistance cassette was cloned between the 1000 base pairs of DNA that flank the 5’ and 3’ ends of *esaA* including the first 15 bases of the *esaA* ORF at the end of 5’ flank and the last 15 bases of *esaA* at the start of the 3’ flank. The final plasmid for allelic replacement was pETduet-1::5’*esaA*flank_SpecPromoter_*kanR*_3’*esaA*flank. This plasmid was then digested with BamHI and NotI and the resulting insert (5’*esaA*flank_SpecPromoter_*kanR*_3’*esaA*flank) was gel extracted (Monarch DNA Gel Extraction Kit, NEB). 100 ng of purified insert was transformed into *S. intermedius* B196 and plated onto THY agar plates supplemented with 250 μg/ml of kanamycin. PCR was used to confirm deletion of *esaA*.

### Secretion assays

Overnight cultures of *S. intermedius* strains were centrifuged at 7600 *g*, resuspended in 1.6 ml fresh THY broth. These washed cultures were then used to inoculate 5 ml THY broth in 15 ml polypropylene centrifuge tubes to an initial OD of 0.1. Cells were harvested (4000 rpm, 4°C, 15 min) when they reached OD_600_ 0.7-0.9 and supernatant fractions were prepared as follows. 3.5 ml of supernatant was removed and filtered through a 0.2 µm membrane to remove remaining cells. Proteins were precipitated at 4°C for 30 minutes by adding 700 µl of cold 100% trichloroacetic acid (TCA, final concentration 16.7%). Precipitant was collected by centrifugation (swinging-bucket, 4600 rpm, 4°C, 30 min), and washed 3 times with 500 ul of cold acetone. Precipitant was then air dried in a fume hood for at least 30 minutes before being dissolved in 20 µl resuspension buffer (50 mM Tris:HCl pH 8.0, 150 mM NaCl, 1X protease cocktail inhibitor). Cell fractions were prepared as follows. Cell pellets were washed with 1 ml PBS, transferred to a 2 ml centrifuge tube, re-pelleted (10,000 *g*, 4°C, 10 min), decanted and snap frozen at -80°C. Washed pellets were then resuspended in 50 μl of lysis buffer (50 mM Tris:HCl pH 8.0, 150 mM NaCl, 10 mg/ml lysozyme, 1X protease cocktail inhibitor), and incubated at 37°C for half an hour. Cell numbers were matched across samples by diluting cells in PBS based on final culture OD_600_. Matched samples were then prepared for western blotting by mixing 2:1 with 4X SDS-PAGE loading dye (125 mM Tris:HCl pH 6.8, 20% *v*/*v* glycerol, 0.01% *w*/*v* bromophenol blue, 4% *v*/*v* BME), heated at 95°C for 10 minutes, and centrifuged (21,000 *g*, room temperature, 15 minutes).

### Antibody generation and western blot analyses

Custom polyclonal antibodies for *S. intermedius* EsaA, EsxA and SodA were generated for this study (Customer’s Antigen Polyclonal Antibody Package, Genscript). C-terminally his_6_-tagged *Si*EsaA_41-871_, EsxA and SodA were purified as described in “Protein purification and expression” except that PBS was used in place of Tris:HCl for all purification buffers. 10 mg of each protein was sent to Genscript for antibody production. Generation of the α-TelC antibody has been described previously (Whitney et al., 2017).

With the exception of EsxA, western blot analyses of protein samples were performed using a Tris-glycine gel and buffer system and a standard western blotting protocol. The SDS-PAGE system for EsxA blots required the use of a tris-tricine buffer system, which allows for the electrophoretic separation of low molecular weight proteins. After SDS-PAGE separation, proteins were wet-transferred to 0.45 µm PVDF membranes (80 V for 1 hour, 4°C). Cell and supernatant fractions were analyzed by Western blot using the protein-specific rabbit primary antibodies α-TelC (1:5000 dilution, 1.5 hours), α-EsaA (1:5000, 1 hour), α-EsxA (1:5000, 2 hours), α-SodA (1:5000, 30 minutes), α-VSV-G (1:3000, 1.5 hours) and a goat α-rabbit secondary antibody (Sigma, 1:5000, 45 minutes). Clarity Max Western ECL substrate (Bio-Rad) was used for chemiluminescent detection of the secondary antibody and all blots were imaged with a ChemiDoc XRS+ System (Bio-Rad).

### NADase activity assay

The consumption of NAD^+^ by *S. intermedius* culture supernatants was assayed as described previously (Whitney et al., 2017). Briefly, culture supernatants taken from mid-log cultures were concentrated 50-fold by spin filtration at 3000 *g* (10kDa MWCO) and then filtered through a 0.2 μm membrane. The samples were then incubated 1:1 with PBS containing 2 mM NAD^+^. Reactions were incubated overnight (approximately 16 hours) at room temperature. 6M NaOH was added to terminate the reaction which was then incubated for 15 minutes in the dark. Fluorescence (ex: 360nm, em: 530nm) was measured using a Synergy 4 Microplate Reader (BioTek Instruments).

### Subcellular fractionation by ultracentrifugation

1L *S. intermedius* cultures were grown to OD_600_ = 0.8 and pelleted by centrifugation at 6000 *g*. Pellets were resuspended in 20 ml of lysis buffer (20 mM Tris:HCl pH 7.5, 150 mM NaCl, 2 mg/ml lysozyme), incubated at 37°C for one hour, and sonicated at 30% amplitude for three pulses of 30 seconds each. The insoluble cellular debris was cleared by centrifugation at 39,191 *g*. The resulting supernatant was then centrifuged for two hours at 200,000 *g* to isolate the membrane fraction. The resulting supernatant (cytosolic fraction) was mixed 1:1 with Laemmli loading buffer. The membrane pellet was washed once with 20 mM Tris:HCl, pH 7.5, 150 mM NaCl, before being resuspended in Laemmli loading buffer. Cytoplasmic and membrane fractions were then analyzed by SDS-PAGE and Western blot.

### Membrane topology mapping

The cysteine labelling experiment was adapted from Ruhe et al. (Ruhe et al., 2018). Briefly, 20 ml cultures of *S. intermedius* strains were grown to OD_600_ = 0.5 and harvested by centrifugation at 4,000 *g* for 20 minutes. Cell pellets were then washed three times with PBS to remove any extracellular material. The pellets were resuspended in 35 μl PBS, pH 7.2 and IRDye680LT-maleimide dye (LI-COR Biosciences) was added to cells to a final concentration of 40 μM. The reactions were incubated at room temperature for 30 minutes in a darkroom before being quenched by adding BME to final concentration of 6 mM. Cells were then harvested by centrifugation and washed three times with PBS supplemented with 6 mM BME. Washed pellets were resuspended in SDS-loading dye and boiled for 10 minutes. Samples were run on SDS-PAGE and imaged with a Chemidoc system (Bio-Rad) using a red LED epi-illumination source and a 700nm/50mm band pass filter.

### Protein expression and purification

*E. coli* BL21(DE3) CodonPlus strains containing pET29b-derived plasmids were grown to OD_600_ = 0.4 and protein expression was induced with 1 mM IPTG. The induced strains were incubated overnight (approximately 18-20 hours) in a 225 rpm shaking incubator at 18°C after which the cells were collected by centrifugation at 10,000 *g*. Cell pellets were resuspended in lysis/wash buffer (20 mM Tris-HCl pH 7.5, 300 mM NaCl, 10 mM imidazole) and sonicated four times at 30% amplitude for 30 seconds each to lyse cells. Cleared cell lysates were purified by affinity chromatography using a Ni-NTA agarose column. After passing cell lysates over the column, the Ni-NTA resin was washed four times using wash buffer and eluted with wash buffer supplemented with 400 mM imidazole. Protein samples were further purified by size exclusion chromatography using a HiLoad 16/600 Superdex 200 column connected to an AKTA protein purification system (Cytiva).

### Crystallization and structure determination

Selenomethionine incorporated *Sg*EsaA_235-829_ was concentrated to 10mg/ml by spin filtration using an Amicon Ultra Centrifugal filter unit with a 30kDa pore size (Millipore). Concentrated protein was screened for crystallization with the MCSG Crystallization Suite (Anatrace). Long, slender crystals formed in 0.2M MgCl_2_, 0.1M Tris:HCl, pH 7.0, 10% (w/v) PEG 8000, after three weeks. Protein crystallization was optimized around this condition with crystals forming in 0.1M Tris:HCl pH 7.0-7.8 and 10-15% (*w*/*v*) PEG 8000. Crystals were cryo-protected in similar buffers supplemented with 20% (*v*/*v*) ethylene glycol. X-ray data were collected at the Structural Biology Center (SBC) sector 19-ID at the Advanced Photon Source. A total of 290 diffraction images of 0.5° for 0.5 sec/image were collected on a Dectris Pilatus3 × 6M detector with a crystal to detector distance of 540 mm. Data were indexed, integrated, and scaled using the *xia2* system (Winter et al., 2013).

The structure of selenomethionine incorporated *Sg*EsaA_235-829_ was solved using the Se-SAD method using the AutoSol package in Phenix (Adams et al., 2010). The AutoBuild wizard was subsequently used for model building and the observed electron density allowed model building for residues 330-727 *Sg*EsaA_235-829_ with an unmodeled gap between residues 514-554 (Terwilliger et al., 2008). Manual adjustments to the model were performed in COOT and model refinement was carried out with Phenix.refine resulting in final *R*_work_ and *R*_free_ values of 0.21 and 0.26, respectively (Afonine et al., 2012; Emsley and Cowtan, 2004). X-ray data collection and refinement statistics are listed in Table 1.

### Homology modeling of *Si*EsaA and PIP

Homology models of *Si*EsaA and *E. faecalis* V583 Phage Infection Protein (PIP) were generated based on our solved structure of *Sg*EsaA_235-829_ using the PHYRE^2^ one-to-one threading algorithm (Kelley et al., 2015). *Si*EsaA was modeled with 100% confidence over 325 residues. *E. faecalis* V583 PIP was modeled with 96% confidence over 341 residues.

### Sequence alignments and conservation mapping

Protein sequence conservation was mapped onto the structure of *Sg*EsaA using the online ConSurf server (Ashkenazy et al., 2016). The multiple sequence alignment used in the calculation was generated as follows. The full-length protein sequence of *Sg*EsaA was used as a BLASTp query sequence against the NCBI Reference Protein Sequence database, restricted to the phylum Firmicutes, using otherwise default settings (Altschul et al., 1990). Full length sequences for the top 500 hits were downloaded and sequences shorter than 750 amino acids were filtered out. A multiple sequence alignment using the remaining 434 sequences was generated using Clustal Omega and uploaded with the structure coordinates (Sievers et al., 2011). Dimer interface calculations for *Sg*EsaA, including buried surface area and ΔG^i^, were performed by uploading structure coordinates to the PDBePISA server (Krissinel and Henrick, 2007).

### SEC-MALS analysis

Size exclusion chromatography with multi-angle laser static light scattering was performed on *Si*EsaA_328-685_ and *Sg*EsaA_332-725_. The proteins were expressed and purified as described above, concentrated to 2 mg/ml by spin filtration and then run on a Superdex 200 column (GE Healthcare). MALS was conducted using a MiniDAWN and Optilab system (Wyatt Technologies). Data was collected and analyzed using the Astra software package (Wyatt Technologies).

### Cysteine crosslinking experiments

For in vitro crosslinking experiments, each cysteine mutant was expressed in *E. coli* BL21 (DE3) CodonPlus and the resulting protein was purified by Ni-NTA affinity chromatography. The eluted protein samples were exposed to environmental oxygen for 16 hours to allow for crosslinking to occur. Samples were then mixed 1:1 with Laemmli buffer either containing or lacking β-mercaptoethanol and analyzed by SDS-PAGE and Coomassie staining. The SDS-PAGE gels were imaged using a ChemiDoc MP system (BioRad).

*In vivo* cysteine crosslinking was conducted similarly except that a pDL277 plasmid-based system was used to express each cysteine mutant in a *S. intermedius* B196 Δ*esaA* background. *S. intermedius* strains were grown to OD_600_ = 0.8 and centrifuged at 4000 *g*. Pellets were resuspended in 200 μl of lysis buffer (20 mM Tris:HCl, pH 7.5, 150 mM NaCl, 1% *w*/*v* DDM, 10 mg/ml lysozyme) and incubated at 37°C for one hour. Samples were then sonicated three times at 30% amplitude for 15 seconds per pulse. Lysed samples were cleared by centrifugation at 21,130 *g* for 20 minutes. The supernatant was then removed and allowed to sit at room temperature for one hour to allow for crosslinking. Samples were analyzed by Western blot using an α-EsaA primary antibody.

### Accession numbers

The atomic coordinates and structure factors for the *Sg*EsaA_235-829_ crystal structure have been deposited in the Protein Data Bank (http://wwpdb.org/) under the PDB code 7JQE.

## Acknowledgements

The authors would like to thank Nathan Bullen, Shehryar Ahmad and Gerd Prehna for constructive feedback on the manuscript and Madoka Akimoto and Giuseppe Melacini for assistance with SEC-MALS experiments. We sincerely thank the members of the SBC at Argonne National Laboratory for their help with data collection at 19-ID beamline. The use of SBC beamlines at the Advanced Photon Source is supported by the U.S. Department of Energy (DOE) Office of Science and operated for the DOE Office of Science by Argonne National Laboratory under Contract No. DE-AC02-06CH11357. T.A.K. is supported by an Alexander Graham Bell Canada Graduate Scholarship from the Natural Sciences and Engineering Research Council of Canada (NSERC). This work was supported by start-up funds from McMaster University.

## Author Contributions

T.A.K. and J.C.W. planned the study. All authors contributed to experimental design. T.A.K. and J.C.W. generated strains and plasmids. T.A.K. performed protein expression, purification and crystallization. S.Y.G. and V.S.S. assisted with protein crystallization. T.A.K., D.W.G., Y.K. and J.C.W. solved and analyzed the crystal structure. T.A.K. and D.W.G. performed biochemical experiments. T.A.K., D.W.G., and J.C.W. analyzed the data. T.A.K., D.W.G. and J.C.W. wrote the paper. All authors provided feedback on the manuscript.

## Supplemental Data

**Supplementary Figure 1.**
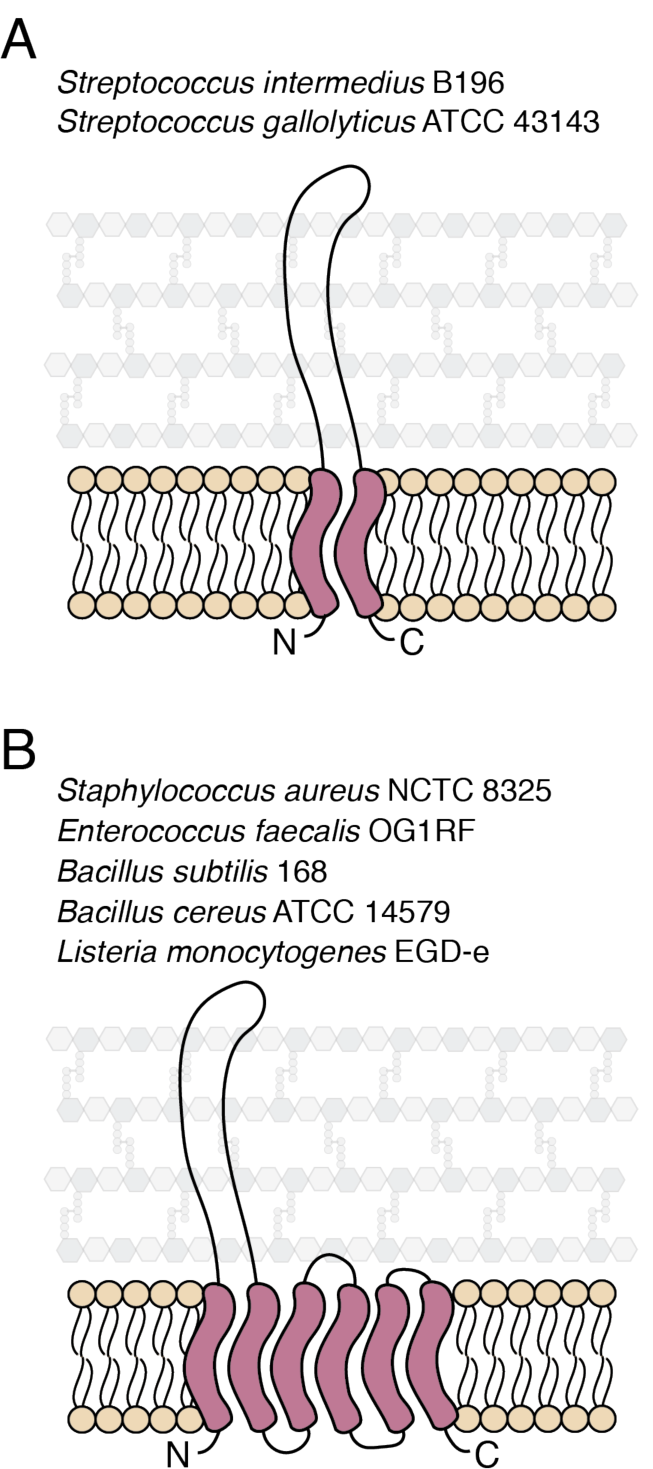
Schematic depicting the two common predicted membrane topologies of EsaA. (A-B) EsaA proteins typically have one N-terminal and one C-terminal TMD (A) or one N-terminal and five C-terminal TMDs (B). TMDs as predicted by TMHMM are depicted in red (Krogh et al., 2001). Several representative strains of Firmicutes bacteria are listed for each topology.

**Supplementary Figure 2.**
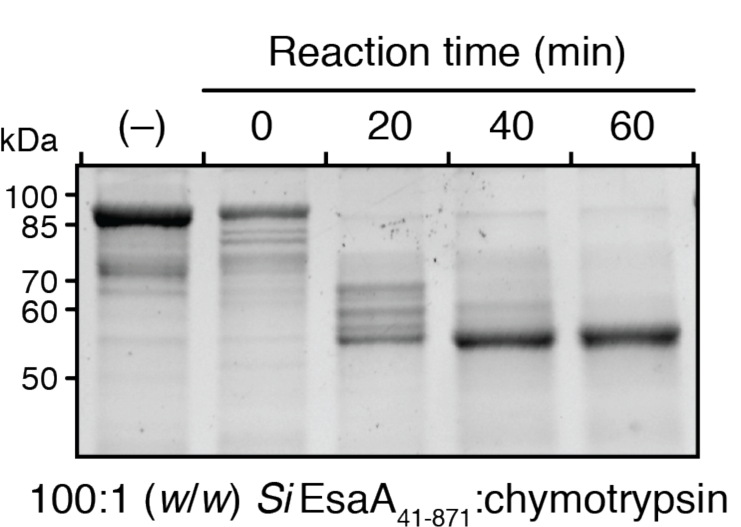
Digestion of *Si*EsaA_41-871_ with chymotrypsin results in a stable truncation of approximately 55kDa. A 1:100 (*w*/*w*) chymotrypsin: *Si*EsaA_41-871_ digestion was conducted over one hour with samples being taken every 20 minutes. The (–) condition indicates untreated *Si*EsaA_41-871_. *Si*EsaA_41-871_ has a predicted molecular weight of 92.5kDa and the amino acid sequence of the 55kDa truncation of *Si*EsaA_41-871_ was confirmed by liquid chromatography-tandem mass spectrometry.

**Supplementary Figure 3.**
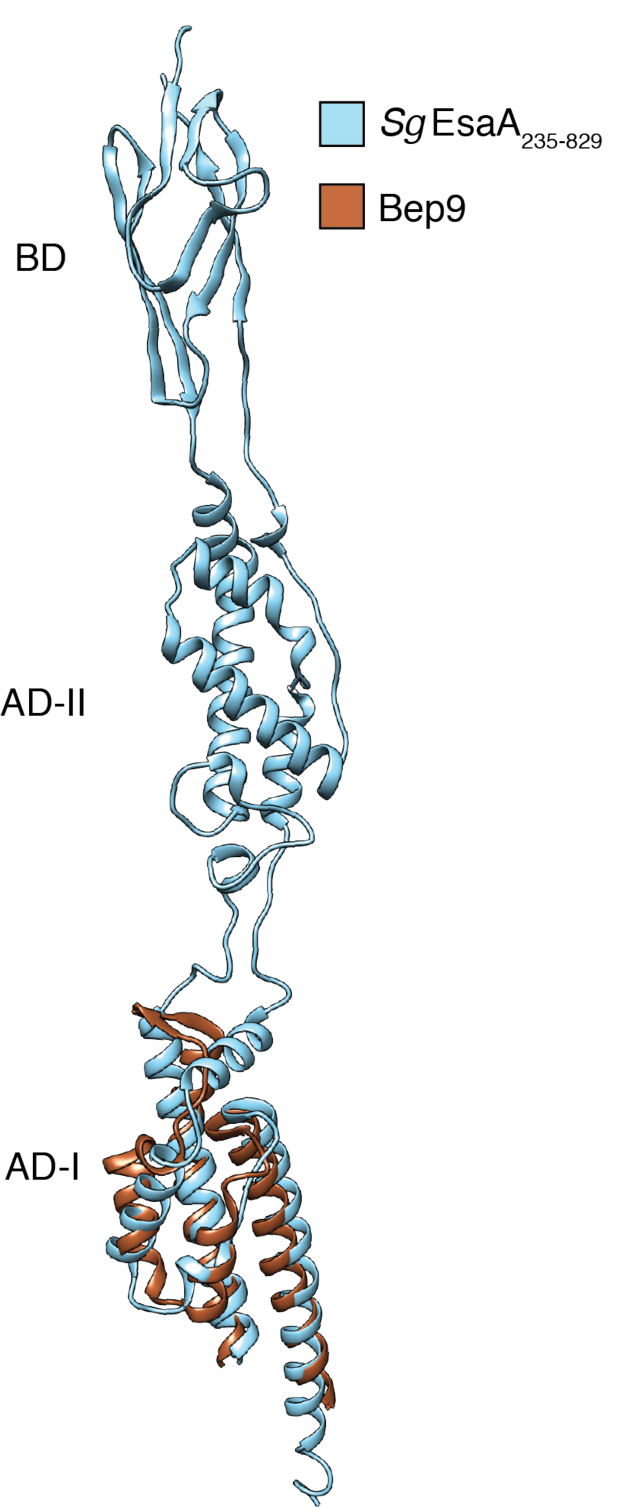
The BID domain of Bep9 from *Bartonella clarridgeiae* resembles the AD-I domain of *Sg*EsaA_235-829_. Bep9 (PDB code 4YK2) is the highest scoring structural homologue for *Sg*EsaA_235-829_ as determined by DALILITE (Z-score, 8.5; Cα root mean squared deviation of 3.5Å over 100 aligned residues). The structures of Bep9 (orange) and *Sg*EsaA_235-829_ (blue) were superimposed using UCSF Chimera.

**Supplementary Figure 4.**
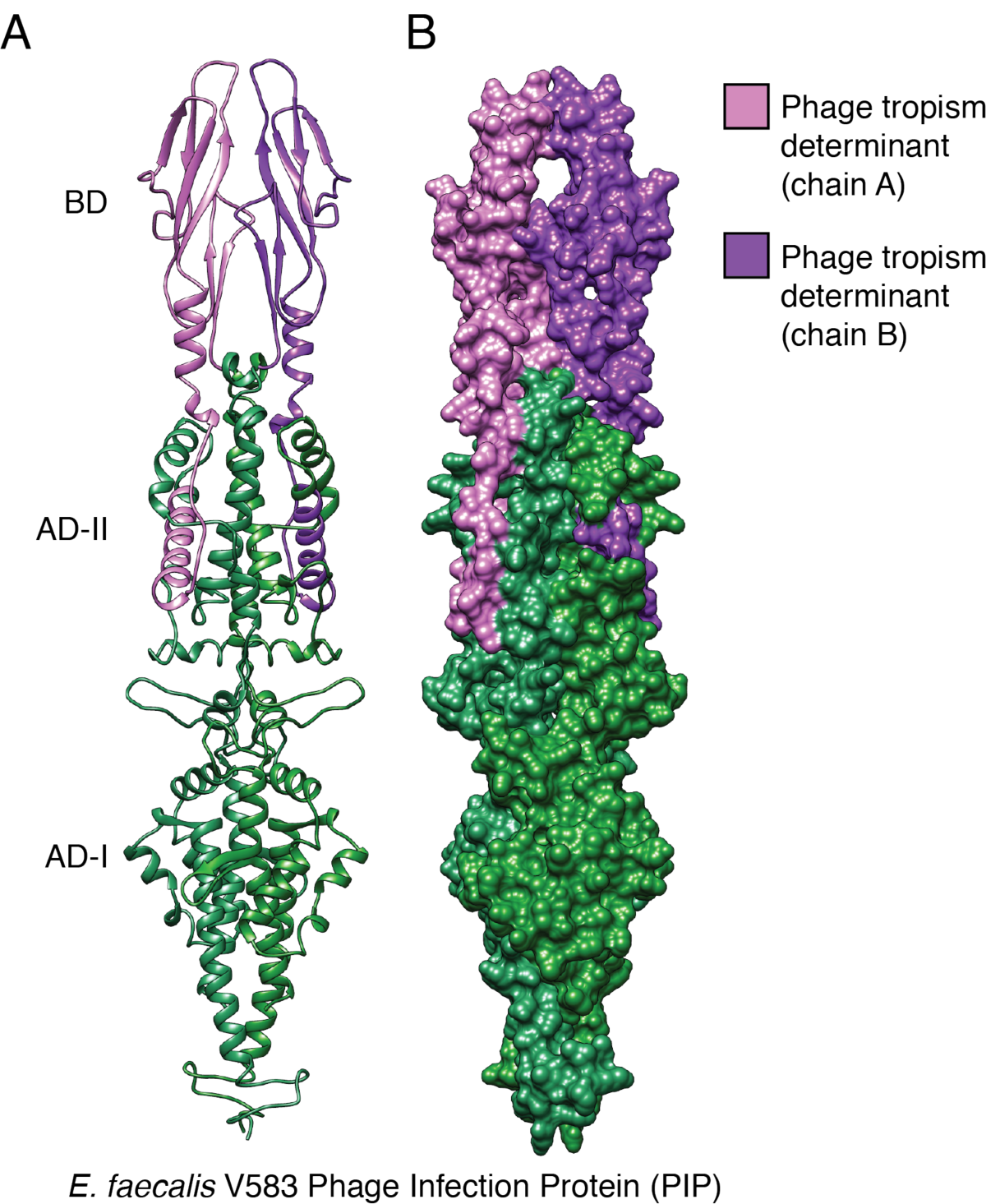
The structure of *Sg*EsaA_235-829_ allows for homology modelling of *E. faecalis* V583 PIP. (A) Ribbon and (B) surface diagrams of the structure of the *E. faecalis* V583 PIP protein were generated by the one-to-one threading algorithm of Phyre^2^ (Kelley et al., 2015). The phage tropism region, coloured pink and purple, encompasses the BDs and a short segment of the AD-II domains of the *Sg*EsaA_235-829_ homodimer.

**Table S1:** Bacterial strains used in this study.

| Organism | Genotype | Description | Reference |
| --- | --- | --- | --- |
| *S. intermedius* B196 | Wild-type |  | (Olson et al., 2013) |
| | ΔSIR_0175::kan<sup>R</sup> | *essC* deletion strain | (Klein et al., 2018) |
| | ΔSIR_0176::kan<sup>R</sup> | *esaA* deletion strain | This study |
| *S. gallolyticus*<br>ATCC 43143 | Wild-type |  |  |
| *E. coli* XL-1 Blue | *recA1 endA1 gyrA96 thi-1*<br>*hsdR17 supE44 relA1*<br>*lac* [F' *proAB*<br>*lacI<sup>q</sup> Z $\Delta$ M15 Tn10* (Tet<sup>R</sup>)] | Cloning strain | Agilent |
| *E. coli* BL21 (DE3)<br>CodonPlus | F<sup>-</sup> *ompT gal dcm lon*<br>*hsdS<sub>B</sub>(r<sub>B</sub><sup>-</sup> m<sub>B</sub><sup>-</sup>)* $\lambda$(DE3)<br>pLysS(Cm<sup>R</sup>) | Protein expression<br>strain | Novagen |
| *E. coli* B834 (DE3) | F<sup>-</sup> *ompT hsdS<sub>B</sub>(r<sub>B</sub><sup>-</sup> m<sub>B</sub><sup>-</sup>)*<br>*$\lambda$(DE3) gal dcm met* | Methionine<br>auxotroph | Novagen |

**Table S2:**
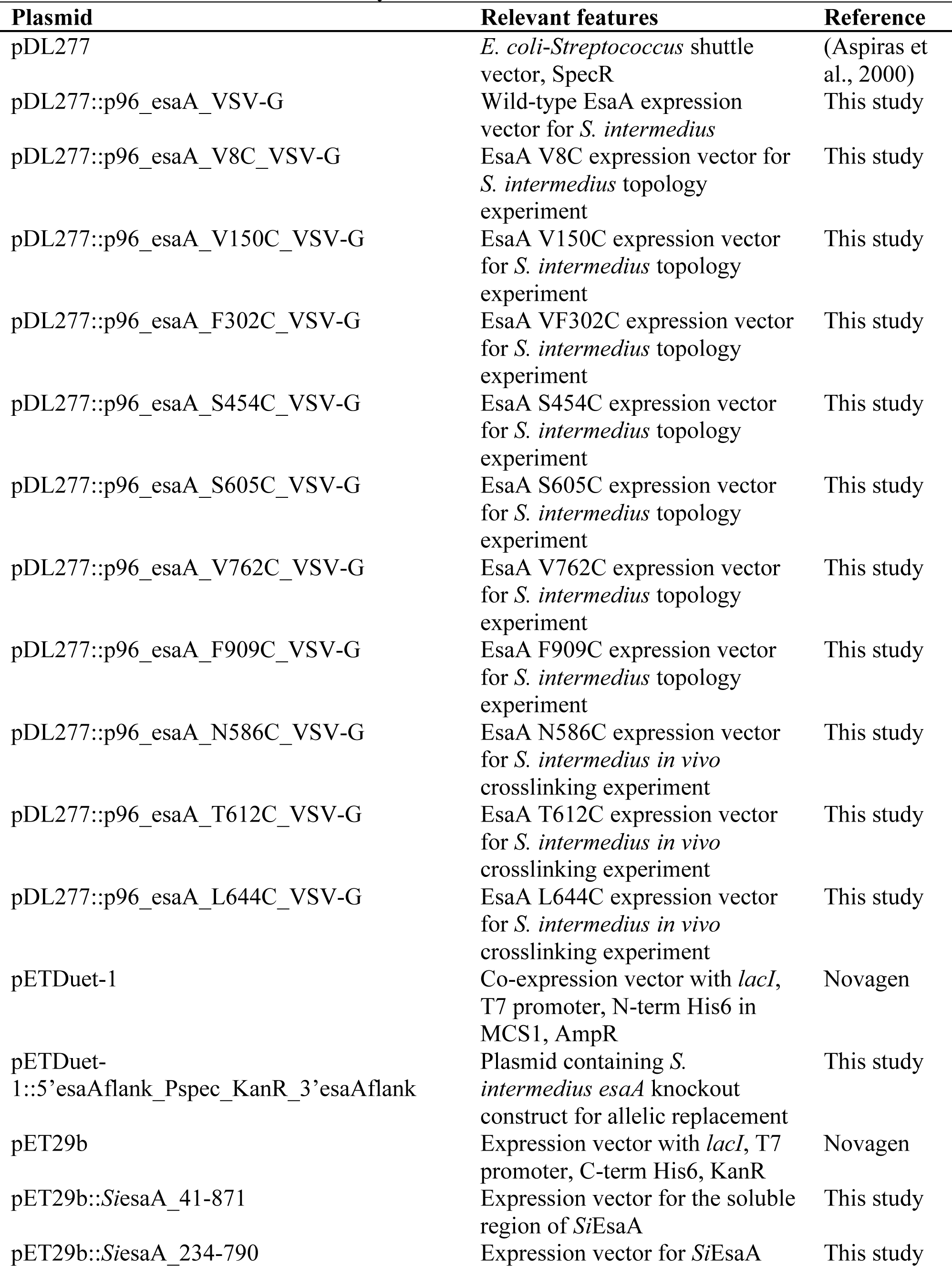
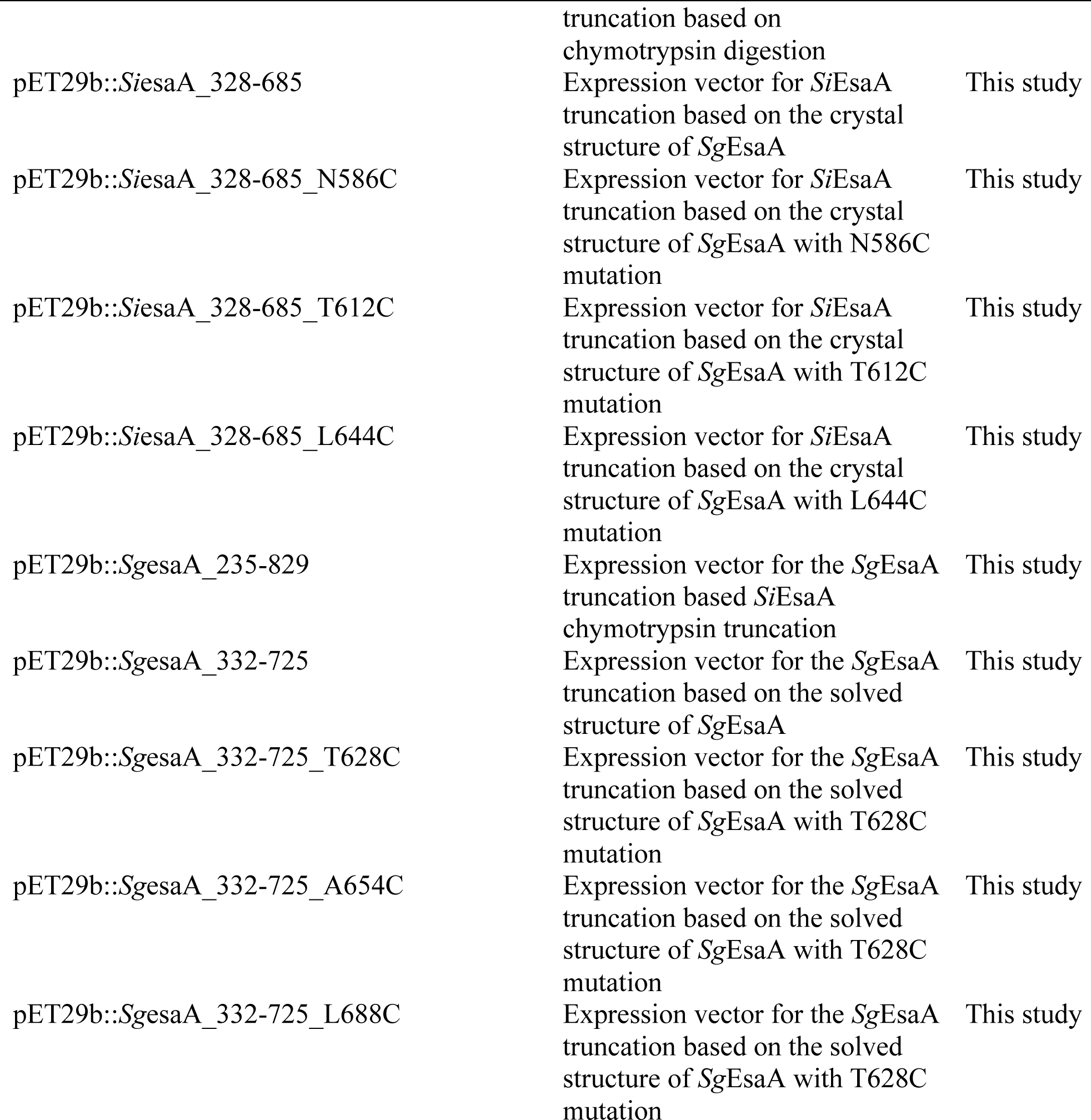
Plasmids used in this study.

